# The virulome of *Streptomyces scabiei* in response to cello-oligosaccharides elicitors

**DOI:** 10.1101/2021.08.10.455888

**Authors:** Benoit Deflandre, Nudzejma Stulanovic, Sören Planckaert, Sinaeda Anderssen, Beatrice Bonometti, Latifa Karim, Wouter Coppieters, Bart Devreese, Sébastien Rigali

**Author notes:** B.D. and N.S. contributed equally to this work. Repositories: RNAseq data were publicly deposited, and our experimental and analytical pipeline were described on the GEO database repository (Accession number: GSE181490).

## Abstract

2.

The development of spots or lesions symptomatic of the common scab disease on root and tuber crops is caused by few pathogenic *Streptomyces* with *Streptomyces scabiei* 87-22 as the model species. Thaxtomin phytotoxins are the primary virulence determinants, mainly acting by impairing cellulose synthesis, and their production in *S*. *scabiei* is in turn boosted by the cello-oligosaccharides released from host plants. In this work we aimed to determine which molecules and which biosynthetic gene clusters (BGCs) of the specialized metabolism of *S. scabiei* 87-22 show a production and/or transcriptional response to cello-oligosaccharides. Comparative metabolomic and transcriptomic analyses revealed that molecules of the virulome of *S. scabiei* induced by cellobiose and cellotriose include i) thaxtomins and concanamycins phytotoxins (and to a lesser extent N-coronafacoyl-L-isoleucine), ii) desferrioxamines, scabichelin and turgichelin siderophores in order to acquire iron essential for housekeeping functions, iii) ectoine for protection against osmotic shock once inside the host, and iv) bottromycins and concanamycins antimicrobials possibly to prevent other microorganisms from colonizing the same niche. Importantly, both cell-oligosaccharides reduced the production of the spore germination inhibitors germicidins and the plant growth regulators rotihibins. The metabolomic study also revealed that cellotriose is in general a more potent elicitor of the virulome compared to cellobiose. This result supports an earlier hypothesis that suggested that the trisaccharide would be the real virulence-triggering factor released from the plant cell wall through the action of thaxtomins. Interestingly, except for thaxtomins, none of these BGCs’ expression seems to be under direct control of the cellulose utilization repressor CebR suggesting the existence of another master regulator sensing the internalization of cello-oligosaccharides. Finally, we found nine additional cryptic and orphan BGCs that have their expression awakened by cello-oligosaccharides, demonstrating that other and yet to be discovered metabolites are part of the virulome of *S*. *scabiei*.

**Impact statement:** Unveiling the environmental triggers that signal proper conditions for host colonization and what is the composition of the arsenal of metabolites specialized for this task (the virulome) is key to understand host-pathogen interactions. In this work, focused on the induction of the common scab disease caused by *Streptomyces* species, we provided further knowledge to both aspects i.e., i) highlighting the capability of cellotriose to trigger the entire virulome and not only the production of thaxtomin phytotoxins, and ii) identifying the set of metabolites that specifically respond to cello-oligosaccharides emanating from the plant under attack. Importantly, we also revealed that the expression of nine cryptic/orphan biosynthetic gene clusters (BGCs) involved in the production of unknown compounds was drastically activated upon cello-oligosaccharides import suggesting that a significant part of the virulome of *S*. *scabiei* remains to be discovered. Finally, we unexpectedly found that the expression control of most of the known and cryptic BGCs does not depend on the cello-oligosaccharide utilization repressor CebR which suggests the existence of another and yet unknown master regulator of the virulence in *S*. *scabiei*.

**Significance as a BioResource to the community:** Not Applicable

**Outcome:** Not Applicable

**Data summary:** [A section describing all supporting external data including the DOI(s) and/or accession numbers(s), and the associated URL.]

The authors confirm all supporting data, code and protocols have been provided within the article or through supplementary data files. RNAseq data were publicly deposited, and our experimental and analytical pipeline were described on the GEO database repository (Accession number: GSE181490)

## 7. Introduction

*Streptomyces scabiei* (syn. *S*. *scabies*) is responsible for causing the disease called “common scab” (CS) on root and tuber crops. Together with a dozen of other phylogenetically related *Streptomyces* species, *S*. *scabiei* colonizes and infects underground storage organs like potato tubers, beets, radishes, turnips, carrots, and peanuts (1, 2). CS lesions cause significant economic losses throughout the world, with potato being the most affected crop. The range of symptoms and lesion morphologies on potato tubers goes from superficial to raised or deep- pitted scabs (3). Although CS is characterized by skin defects, it mostly affects the visual aspect of the tuber tissues and root, sometimes also reducing their size, causing a significant drop in the quality and marketability of the potato tubers (3).

The virulence factors which predominantly contribute to the development of CS are the thaxtomin phytotoxins which are nitrated diketopiperazines (4). Eleven thaxtomin analogues have been identified up to date, thaxtomin A being the predominant form associated with the disease (5–7). While the molecular targets are still unknown, thaxtomin A alters the expression of host genes involved in cellulose biosynthesis, cell wall remodeling and strengthening (8, 9). Cellulose synthase complexes were also shown to be affected in their density and motility (8), possibly due to endocytosis triggered by thaxtomin A (10). *In vivo*, thaxtomin A causes multiple symptoms to plant targets (11)including necrosis, perturbation of ion fluxes, cell hypertrophy, callose deposition, and ectopic lignin formation. In addition, wounded or immature sites are affected, inducing synthesis of hemicellulose and pectins and leading to the deposition and excessive accumulation of layers of periderm (12).

Next to thaxtomins, other specialized metabolites are known or predicted to play important roles in plant colonization and infection by *S. scabiei*. Several studies revealed that concanamycins and coronafacoyl phytotoxins also contribute to the development of plant disease (11). In synergy with thaxtomins, concanamycins were shown to play an essential role in the type and morphology of developed lesions (13). Contrary to concanamycins and thaxtomins, the exact roles of coronafacoyl phytotoxins in CS disease development are still relatively vague and were shown to be non-essential for pathogenicity development (11). However, their impact via modulating jasmonate hormone signalling networks could assist in overcoming host defence mechanisms (14). N-coronafacoyl-l-isoleucine (CFA-Ile) is the major product of the coronafacoyl gene cluster (15). A wide spectrum of virulence-associated activities of CFA-Ile, like tissue hypertrophy, leaf chlorosis and inhibition of root elongation, were reported in plants. Nevertheless, coronafacoyl phytotoxins are found in non-pathogenic *Streptomyces* spp. as well, suggesting some additional unidentified roles along with the already cited disease-related activities (16). Recently, the production of two novel phytotoxic metabolites was highlighted in *S. scabiei*. Rotihibins C and D are lipopeptides that significantly reduce the photochemistry efficiency of the photosystem II which in turn affects the growth of *Arabidopsis thaliana* and *Lemna minor* at low concentrations. At even lower concentrations, *L. minor* plantlets exhibit an increase in their surface area, suggesting a hormetic effect of rotihibins (17).

Siderophores are also key metabolites for host infecting bacteria, iron being indispensable for housekeeping functions such as DNA replication and protein synthesis. Next to desferrioxamines that are essential for *Streptomyces* survival in iron limited environments (18), S. scabiei and related species have the ability to produce diverse and specific siderophores including scabichelin and turgichelin (19), as well as pyochelin (20) together with three other yet unknown iron chelators deduced from genome mining analysis. This multitude of siderophores with high affinity for ferric iron would guarantee *S. scabiei* to capture iron trapped in its hosts. However, although iron acquisition might contribute to the onset of pathogenicity, *in planta* bioassays showed that there is no connection between virulence and pyochelin production by *S. scabiei* (20).

How *S. scabiei* senses the presence of its plant host and triggers its specialized metabolism required for virulence has also been a main research topic. Induction of thaxtomin biosynthesis requires the import of cello-oligosaccharides (cellobiose and/or cellotriose) by the sugar ABC transporter composed by CebE as the sugar-binding component, CebF and CebG as components of the membrane permease, and MsiK to provide energy to the transport via ATP hydrolysis (21–23). Once inside the cytoplasm, the imported cello-oligosaccharides inhibit the DNA-binding ability of the transcriptional repressor CebR which in turn allows the expression of the *txt* cluster pathway-specific activator TxtR (22, 24).

If the path from cello-oligosaccharide uptake to activation of thaxtomin biosynthesis is well described at the molecular level, many questions remain unsolved. The first issue regards cellobiose itself as a natural environmental elicitor of the CS disease. Most *in vivo* and *in vitro* studies on the induction of the pathogenic lifestyle of *S*. *scabiei* used cellobiose and not cellotriose as the triggering factor. The only reason why cellotriose is usually excluded from laboratory studies is because it is much more expensive and less available in large quantities compared to cellobiose. However, earlier studies suggested that cellobiose is unlikely to be released from plant cell wall depolymerization by thaxtomins during host colonization by *S*. *scabiei*. Instead, incubation of tobacco and radish seeds with thaxtomin A showed release of cellotriose (25). Cellobiose on the other hand, is an important by-product of cellulose hydrolysis by the cellulolytic system, which naturally occurs upon organic matter turnover by the soil microflora. However, the ability of *S*. *scabiei* to degrade cellulose is insignificant despite possessing a complete cellulolytic system (25–27). While the molecular mechanism silencing the cellulolytic system of *S*. *scabiei* is unknown, it avoids the release of cellobiose from decaying plant biomass, hence preventing this bacterium to be a protagonist in the mineralization of organic soils. This particularity could be a major evolving adaptation that somehow “forces” *S*. *scabiei* to instead colonize living plant tissues. Sensing cellotriose released by the depolymerization of cellulose caused by thaxtomin, together with silencing the cellulolytic system and thus avoiding the release of cellobiose, could allow this bacterium to discriminate if cellulose by-products originate from living or instead dead plant cell walls (25, 27).

An even more important question that remains to be solved is the exact composition of the arsenal of specialized metabolites that constitute the virulome of *S*. *scabiei*. Are thaxtomins the only phytotoxins that respond to virulence elicitors or do other specialized metabolites display the same production response? Recently, the group of Prof Dawn Bignell has shown that cultivation of *S*. *scabiei* in the oat bran agar (OBA) medium is not only able to induce thaxtomin production, but also other specialized metabolites known or predicted to play an important role in colonizing and infecting the plant host tissues, i.e., CFA-Ile, concanamycins, siderophores (desferrioxamines and pyochelin), and also a form of auxin (IAA) (28). OBA is a complex plant-based medium in which cello-oligosaccharides are proposed to be responsible for the induction of thaxtomin production (25). However, it cannot be excluded that some of the other compounds present in OBA influence – positively or negatively, alone or in combination – the production of specialized metabolites.

As previous studies suggested cellobiose and/or cellotriose to be natural elicitors of the pathogenic response of *S*. *scabiei* (27), their specific contribution to the induction of the metabolome requires further investigation. In this work we wanted to provide answers to whether cellotriose could – equally to cellobiose – trigger the “virulome” of *S*. *scabiei* and if the cello-oligosaccharide mediated induction takes place at the transcriptional level. Our work revealed that cellotriose is a better inducer of the virulome of *S*. *scabiei* compared to cellobiose. Our transcriptomic analysis also shows that cryptic/orphan biosynthetic gene clusters have their expression awakened by cello-oligosaccharides suggesting that yet unknown metabolites would be part of the virulome of *S*. *scabiei*.

## 8. Methods

### Strain and culture conditions

*Streptomyces scabiei* 87-22 and its Δ*cebR* mutant (22) were routinely cultured in Tryptic Soy broth (TSB, 30 g/l, Sigma-Aldrich) or ISP2 (for 1l: 4 g Yeast Extract, 10 g Malt Extract, 4 g Dextrose, pH 7.2) liquid media at 28°C under shaking (180 rpm, New Brunswick™ Innova® 44 incubator shaker). Modified thaxtomin defined medium (TDM) ((25), without L-Sorbose) was used as minimal medium, supplemented with maltose 0.5% (Sigma-Aldrich) in which cellobiose (Carbosynth) and cellotriose (Megazyme) were added as inducers. Sucrose (saccharose) was purchased from Merck.

### Transcriptomics

#### Cultures and sampling

Pre-cultures of *S. scabiei* 87-22 (WT) and Δ*cebR* were conducted in 50 ml ISP2 medium inoculated with 4*10^7^ spores for 24 hours. The mycelium was collected by centrifugation (3,500 g for 5 minutes at room temperature (RT)) and washed twice by resuspension/centrifugation with 20 ml TDM medium without carbon source. The mycelium was then resuspended in TDM + maltose 0.5% (TDMm) or ISP2 to a density of 16 mg/ml (wet biomass) and then split into three Erlenmeyer flasks (per strain and culture condition) containing a culture volume of 25 ml. After 30 minutes of incubation in TDMm at 28 °C, a first sampling (= time points 0) of 2.5 ml was collected from each flask and cellobiose or cellotriose were added to a final concentration of 2.5 mM, each into three flasks. The next samplings were collected following the same procedure, 1 and 2 hours (= time points 1 and 2, respectively) post addition of cello-oligosaccharides. In the ISP2 culture condition involving *S. scabiei* WT and Δ*cebR*, 2.5-ml samples were collected in each flask after 3 hours of culture. All samples were collected in 15-ml Falcon tubes and centrifuged for 3 minutes at 3,500 g (RT). The supernatant was quickly and thoroughly removed, and the tubes were immediately flash-frozen into liquid nitrogen. The frozen cell pellets were placed into a -80°C freezer until RNA extraction.

#### RNA preparation

The RiboPure™ Bacteria RNA Purification Kit (Invitrogen) was used for total RNA extraction. The RNAwiz lysis buffer was added to the frozen mycelium pellets and the procedure was followed according to the manufacturer’s guidelines except the bead-beating step that was extended to 20 minutes. The quantification and quality control of total RNA samples were performed on a Bioanalyzer 2100 (Agilent). Ribosomal RNA depletion and library preparation were carried out using the Ovation Complete Prokariotic RNAseq kit (NuGEN). The libraries were sequenced on a NextSeq® 500 System (Illumina) HM 2X75 bp read length with 7 million reads per library.

#### Read mapping and differential expression

Sequenced reads were quality-checked and trimmed when necessary, using the Trimmomatic Software (29). Reads were subsequently mapped to the reference genome (*S. scabiei 87-22*), using Bowtie2 (30, 31), and an average of 98.7% of reads were aligned. For each transcript, the number of mapped reads were compiled with featureCounts (32), outputting a count table on which the rest of the analysis is based. Differential expression analysis was performed in R, with the DESeq2 package (33). RNAseq data were publicly deposited, and our experimental and analytical pipeline were described on the GEO database repository (Accession number: GSE181490)

#### Metabolomics

After 45 hours of pre-culture in TSB inoculated with 2*10^7^ spores of *S*. *scabiei* 87-22, the mycelium was collected by centrifugation (3,500 g for 5 minutes at RT) and washed twice by resuspension/centrifugation with 20 ml TDM medium without carbon source. The washed mycelium was resuspended to a density of 200 mg/ml (wet biomass) and 1 ml was used to inoculate TDM + maltose 0.5 % (TDMm) plates (25 ml) as overlay. Four conditions were tested with three biological replicates: TDMm; TDMm + cellobiose 2.5 mM; TDMm + cellotriose 2.5 mM; TDMm + sucrose 2.5 mM. After 96 hours of incubation at 28 °C, one half of each plate was extracted with mQ H_2_O (v/v), dried, resuspended in 1 ml of mQ H_2_O, and filtered through 0.22 µm syringe-driven filters. These metabolic extracts were diluted 20 times in 97/3/0.1 H_2_O/ACN/HCOOH to improve chromatographic and mass spectrometric performance.

The µ LC-MS/MS system consisted of a Waters NanoAcquity M-Class UPLC coupled to a Waters Xevo TQ-S triple quadrupole mass spectrometer fitted with an IonKey/MS^TM^ source (Waters, MA, USA). Mobile phase A was 0.1% formic acid in H_2_O (Biosolve) and mobile phase B was 0.1% formic acid in acetonitrile (Biosolve). The strong and weak solutions used to wash the auto-sampler were 0.1% HCOOH in H_2_O and 0.1% HCOOH in acetonitrile/water/isopropanol (Biosolve) (50:25:25, v/v/v), respectively. The samples were directly injected (5 μl injection volume) to a Waters 150 μm x 100 mm, 1.8 μm HSS T3, iKey^TM^ separation device. The metabolites were eluted from the analytical column using the following gradient: 0-10 min: 3-50% B, 10-11 min: 50-80% B, 11-15 min: 80% B, 15-16 min 80-3% B, 16-25 min: 3% B at a flow rate of 2 µl/min. The column was operated at 45°C, ionization was performed in positive mode using a voltage of 3.65 kV. The cone and collision voltage were set respectively at 35 V and 30 V, and the source temperature was 120°C.

Detection was obtained by MRM mode with transitions of the analytes of interest and their specific retention window (± 0.5 min). Selection of these transitions was based on information in the GNPS public spectral library, literature survey, optimization experiments and own findings (17) (Table S1). Data acquisition was performed by MassLynx 4.2 software, and the data was subjected to a Savitzky-Golay smoothing in Skyline v21 (Adams et al. 2020). The Area Under the Curve (AUC) of ion peaks was calculated and normalized to the TDMm condition for each metabolite. Each complete set of different conditions/biological replicates were randomly analysed and separately repeated (triplicates).

#### Genome mining

AntiSMASH (antibiotics and secondary metabolites analysis shell; version 5.1.2), available at https://antismash.secondarymetabolites.org, was used for genome mining (34) in combination with the internal MIBiG 2.0 (Minimum Information about a Biosynthetic Gene cluster) database (35). The complete genome sequence of *Streptomyces scabiei* 87-22 (Ref NC_013929) was used for the prediction of BGCs. Manual inspection was carried out to rectify the synteny values provided by AntiSMASH, only considering protein sequences sharing a minimum of 60% of identity on at least 70% of sequence coverage. BGC delimitation and/or attribution issues were manually corrected and supported by literature survey.

## 9. Results

### The specialized metabolism of *S. scabiei* 87-22

Prior to assessing the transcriptomic and metabolomic responses of *S. scabiei* 87-22 to virulence elicitors, we updated the current knowledge on the BGCs of the specialized metabolism of this species. A genome mining analysis has recently been performed by Liu et al. (28), identifying 34 BGCs including eight terpenes, six non-ribosomal peptide synthetases (NRPSs), six polyketide synthases (PKSs), one hybrid PKS-NRPS BGC, five ribosomally synthesized and post-translationally modified peptides (RiPPs), four siderophores, and four other types of BGC (betalactone, butyrolactone, melanin, and ectoine). We performed additional and manual rounds of inspection (additional BLAST searches and a literature survey) in order to i) identify possible BGC delimitation issues and correct BGC length, ii) split individual BGCs into multiple BGCs, and iii) identify BGCs involved in the production of known natural products absent from the MIBiG database (version 2.0). In total, 12 other BGCs were identified through these additional steps leading to a final list of 46 BGCs (Table 1). Among these 12 additional BGCs, there was only one BGC for which the natural product is known, namely BGC#33b coding for melanin. The other 11 BGCs, including BGC#1b (NRPS), BGC#6b (terpene), BGC#7b (bacteriocin), BGC#16b (lanthipeptide), BGC#23b (butyrolactone), BGC#23c (LAP), BGC#23d (PKS), BGC#27a (Type 1 PKS), BGC#29a (Type 3 PKS) and BGC#31b (linaridin) display relatively low similarity levels with genes of BGCs associated with the biosynthesis of known compounds (Table 1).

**Table 1.** Prediction of BGCs involved in specialized metabolite production in Streptomyces scabiei 87-22

| BGC | BGC genes<br>[old locus tag] | BGC<br>length<br>(pb) | Product type | Specialized<br>metabolite | Bioactivity | Most similar BGC (%)<br>Species | Ref / MIBig ID |
| --- | --- | --- | --- | --- | --- | --- | --- |
| 1a | SCAB_RS00610-00670<br>[SCAB_1361-1481] | 26675 | Siderophore | Pyochelin | Iron uptake | Pyochelin (100)<br>*S. scabiei* 87-22 | (20) |
| 1b | SCAB_RS00675-00760<br>[SCAB_1491-1671] | 18891 | NRPS | Cryptic | Unknown | None | NA |
| 2 | SCAB_RS00870-01005<br>[SCAB_1951-2231] | 34118 | Betalactone | Cryptic | Unknown | Esmeraldin (4)<br>*S. antibioticus* | BGC0000935 |
| 3 | SCAB_RS01465-01545<br>[SCAB_3221-3351] | 32892 | NRPS | Rothibins | Plant growth<br>inhibitory effect | Rothibins (100)<br>*S. scabiei* RL-34 | (17) |
| 4 | SCAB_RS01655-01700<br>[SCAB_3601-3671] | 15765 | Lanthipeptide | Cryptic | Unknown | None | NA |
| 5 | SCAB_RS02290-02370<br>[SCAB_4951-5131] | 19986 | Terpene | 2-methylisoborneol | Smell of soil | 2-methylisoborneol (100)<br>*S. griseus* | (51) |
| 6a | SCAB_RS02505-02545<br>[SCAB_5421-5511] | 11090 | Terpene | Isorenieratene | Light harvesting<br>photoprotection | Isorenieratene (100)<br>*S. argillaceus* | BGC0001456 |
| 6b | SCAB_RS02550-02590 | 7346 | Terpene | Cryptic | Unknown | Guadinomine (4) | BGC0000998 |
|  | [SCAB_5521-5601] |  |  |  |  | *S. sp.* K01-0509 |  |
| 7a | SCAB_RS04050-04080<br>[SCAB_8601-8661] | 13988 | Lanthipeptide | Informatipeptin | Antimicrobial | Informatipeptin (63)<br>*S. viridochromogenes* DSM<br>40736 | BGC0000518 |
| 7b | SCAB_RS04085-04095<br>[SCAB_8681-8701] | 3358 | Bacteriocin | Cryptic | Unknown | None | NA |
| 8 | SCAB_RS05720-05745<br>[SCAB_12041-12091] | 6531 | Butyrolactone | Cryptic | Unknown | Pyocyanine (14)<br>*P. aeruginosa* PA01 | BGC0000936 |
| 9 | SCAB_RS06125-06180<br>[SCAB_12881-13001] | 13913 | Terpene | Hopene | Protection against<br>water loss | Hopene (92)<br>*S. coelicolor* A3(2) | (52) |
| 10 | SCAB_RS08670-08720<br>[SCAB_18341-18441] | 11906 | Siderophore | Cryptic | Unknown | Grincamycin (9)<br>*S. lusitanus* | BGC0000229 |
| 11a | SCAB_RS09255-09340<br>[SCAB_19561-19741] | 27673 | NRPS-like | Cryptic | Unknown | Stenothricin (11)<br>*S. filamentosus* NRRL 15998 | BGC0000431 |
| 11b | SCAB_RS09350-09410<br>[SCAB_19761-19891] | 11446 | NRPS-like | Cryptic | Unknown | s56-p1 (43)<br>*S. cp.* Soc090715ln-17 | BGC0001764 |
| 12 | SCAB_RS09510<br>[SCAB_20121] | 2207 | Terpene | Geosmin | Earthy odorant | Geosmin (100)<br>*S. coelicolor* A3(2) | (53) |
| 13 | SCAB_RS09780-09830<br>[SCAB_20701-20801] | 10413 | Bacteriocin | Cryptic | Unknown | None | NA |
| 14 | SCAB_RS10905-10995<br>[SCAB_23071-23271] | 20828 | Terpene | Cryptic | Unknown | FD-594 (7)<br>*S. sp. Ta-0256* | BGC0000222 |
| 15 | SCAB_RS11635-11660<br>[SCAB_24651-24711] | 9975 | Siderophore | Cryptic | Unknown | None | NA |
| 16a | SCAB_RS15070-15100<br>[SCAB_31761-31841] | 18264 | NRPS | Thaxtomins | Phytotoxin | Thaxtomin A (100)<br>*S. scabiei* 87-22 | (54) |
| 16b | SCAB_RS15145-15225<br>[SCAB_31961-32131] | 19409 | Lanthipeptide | Cryptic | Unknown | None | NA |
| 17 | SCAB_RS20585-20630<br>[SCAB_43271-43361] | 9782 | Type 2 PKS | Spore pigment | Pigment | Spore pigment (75)<br>*S. avermitilis* | BGC0000271 |
| 18 | SCAB_RS20845-20995<br>[NA-44151] | 42317 | NRPS, Type 1 PKS | Cryptic | Unknown | None | NA |
| 19 | SCAB_RS22630-22710<br>[SCAB_47531-47711] | 21090 | Lanthipeptide | Cryptic | Unknown | None | NA |
| 20 | SCAB_RS26995-27050<br>[SCAB_56591-56711] | 17166 | Bacteriocin,<br>bottromycin | Bottromycins | Antibacterial | Bottromycin A2 (100)<br>*S. scabiei* 87-22 | (55) |
| 21 | SCAB_RS27660-27675<br>[SCAB_57921-57951] | 5032 | Siderophore | Desferrioxamines | Iron uptake | Desferrioxamines (100)<br>*S. sp. Id38640* | BGC0001478 |
| 22 | SCAB_RS28265-28270<br>[SCAB_59231-59241] | 1290 | Melanin | Melanin | Pigment | Melanin (100)<br>*S. griseus* | (56) |
| 23a | SCAB_RS30025-30125<br>[SCAB_62881-63081] | 30803 | Type 1 PKS | Cryptic | Unknown | None | NA |
| 23b | SCAB_RS30085-30160<br>[NA-63151] | 23408 | Butyrolactone | Cryptic | Unknown | None | NA |
| 23c | SCAB_RS30120-30205<br>[NA-63271] | 24765 | LAP | Cryptic | Unknown | None | NA |
| 23d | SCAB_RS30125-30265<br>[SCAB_63081-63401] | 37334 | PKS-like | Cryptic | Unknown | None | NA |
| 24 | SCAB_RS33835-33850<br>[SCAB_70711-70741] | 3166 | Ectoine | Ectoine | Osmoprotectant | Ectoine (100)<br>*S. scabiei* 87-22 | (57) |
| 25 | SCAB_RS34860-34975<br>[SCAB_72851-73081] | 24960 | NRPS-like | Cryptic | Unknown | None | NA |
| 26 | SCAB_RS35245-35325<br>[SCAB_73651-73801] | 17977 | Terpene | Cryptic | Unknown | None | NA |
| 27a | SCAB_RS37770-37840<br>[SCAB_78881-79041] | 23685 | Type 1 PKS | Cryptic | Unknown | None | NA |
| 27b | SCAB_RS37860-37955<br>[SCAB_79081-79081] | 27749 | Indole | Cryptic | Unknown | 5-isoprenylindole-3-carboxylate β-D-glycosyl<br>ester (14) | BGC0001483 |
|  |  |  |  |  |  | *S. Sp.* RM-5-8 |  |
| 28 | SCAB_RS38095-38165<br>[SCAB_79581-79721] | 31375 | Type 1 PKS | Coronafacoyl<br>phytotoxins | Phytotoxin | Coronafacoyl phytotoxin,<br>(100)<br>*S. scabiei* 87-22 | (58) |
| 29a | SCAB_RS38310-38370<br>[SCAB_80021-80131] | 15153 | Type 3 PKS | Cryptic | Unknown | Daptomycin (11)<br>*S. filamentosus* NRRL 11379 | BGC0000336 |
| 29b | SCAB_RS38390<br>[SCAB_80171] | 1184 | Type 3 PKS | Germicidin | Inhibitor of<br>germination | Germicidin (100)<br>*S. scabiei* 87-22 | (59) |
| 30 | SCAB_RS39275-39320<br>[SCAB_82111-NA] | 14262 | Terpene | Cryptic | Unknown | None | NA |
| 31a | SCAB_RS40100-40220<br>[SCAB_83841-84101] | 94968 | Type 1 PKS | Concanamycins | Cytotoxic<br>(antifungal,<br>antineoplastic,<br>anti- protozoal<br>and antiviral) | Concanamycin A (89)<br>*S. neyagawaensis* | (49,60) |
| 31b | SCAB_RS40225-40290<br>[NA-84261] | 13911 | Linaridin | Cryptic | Unknown | None | NA |
| 32 | SCAB_RS40385-40435<br>[SCAB_84461-84561] | 13579 | Siderophore | Cryptic | Unknown | None | NA |
| 33a | SCAB_RS40855-40900<br>[SCAB_85431-85521] | 30019 | NRPS | Scabichelin | Iron uptake | Scabichelin (100)<br>*S. scabiei* 87-22 | (19) |
| 33b | SCAB_RS40955-41010<br>[SCAB_85631-85741] | 9584 | Melanin | Melanin | Pigment | Melanin (57)<br>*S. avermitilis* | BGC0000908 |
| 34 | SCAB_RS41165-41255<br>[SCAB_86081-86261] | 19524 | Terpene | Cryptic | Unknown | None | NA |

Over a third of the predicted BGCs (18 out of 46) encode genes involved in the production of natural products that have already been identified in S. *scabiei* or in other *Streptomyces* species (known BGCs) (Table 1). Half of these (9 out of 18) belong to the so-called core metabolome (36) i.e., BGCs involved in the biosynthesis of molecules produced by almost all *Streptomyces* species including 2-methylisoborneol (BGC#5), isorenieratene (BGC#6a), hopene (BGC#9), geosmin (BGC#12), the WhiE spore pigment (BGC#17), desferrioxamines (BGC#21), melanins (BGC#22, BGC#33b), and ectoine (BGC#24). The remaining 9 BGCs of known metabolites were classified into three different functional categories, namely, i) plant-associated molecules (thaxtomins, the coronafacoyl phytotoxins, concanamycins, and rotihibins), ii) siderophores (pyochelin, scabichelin and turgichelin, in addition to desferrioxamines), and iii) antimicrobials (informatipeptin, bottromycins, and germicidins).

Note that concanamycins also exhibit antiviral (37) and antifungal (38) activities due to their capacity to inhibit the V-type H^+^ ATPase (39) and therefore could also have been included into the “antimicrobials” functional category.

The remaining 28 BGCs are considered as “cryptic” or “orphan”, i.e., either their product is a yet undiscovered natural product (unknown unknowns) or is a known compound but the genetic material responsible for its synthesis is still unknown (unknown knowns) (40). Nevertheless, different types of natural products deduced by antiSMASH allowed us to sort these BGCs into different categories. Genome mining predicted four NRPSs (BGC#1b, #11a, #11b, #25), four terpenes (BGC#6b, #14, #26, #30), four PKSs (BGC#23a, #23d, #27a, #29a), three siderophores (BGC#10, #15, #32), three lanthipeptides (BGC#4, #16b, #19), two bacteriocins (BGC#7b, #13), two butyrolactones (BGC#8, #23b), one LAP (BGC#23c), one indole (BGC#27b), one linaridin (BGC#31b), one betalactone (BGC#2), and one hybrid PKS-NRPS BGC (BGC#18) (Table 1).

### Specialized metabolite production upon sensing cellobiose and cellotriose

The effect of cello-oligosaccharides cellobiose and cellotriose on the induction of the specialized metabolism of *S. scabiei* 87-22 was assessed by targeted liquid chromatography-multiple reaction monitoring MS (LC-MRM-MS). Figure 1 shows the Area Under the Curve (AUC) of ion peaks related to known metabolites produced by *S. scabiei* 87-22 (Table S1) when cultured with or without the environmental virulence elicitors cellobiose and cellotriose. Expectedly, thaxtomin A was about 5- and 7-fold overproduced in TDMm + cellobiose and in TDMm + cellotriose, respectively. The greatest production of thaxtomin A appeared to occur upon culture in the presence of cellotriose compared to cellobiose, confirming earlier results suggesting that the trisaccharide has a higher triggering effect on thaxtomin phytotoxin biosynthesis (25). Concanamycins A and B followed the same production pattern with stronger induction rates: on average 16- and 30-fold increases in metabolite levels were observed in TDMm + cellobiose and in TDMm + cellotriose, respectively (Figure 1). In contrast, N-coronafacoyl-L-isoleucine (CFA-Ile) was not strikingly overproduced compared to other plant-associated metabolites. Indeed, we only observed about 2-fold overproduction in both conditions where cello-oligosaccharides were added, and the condition including sucrose displayed similar CFA-Ile levels (Figure 1).

**Figure 1.**
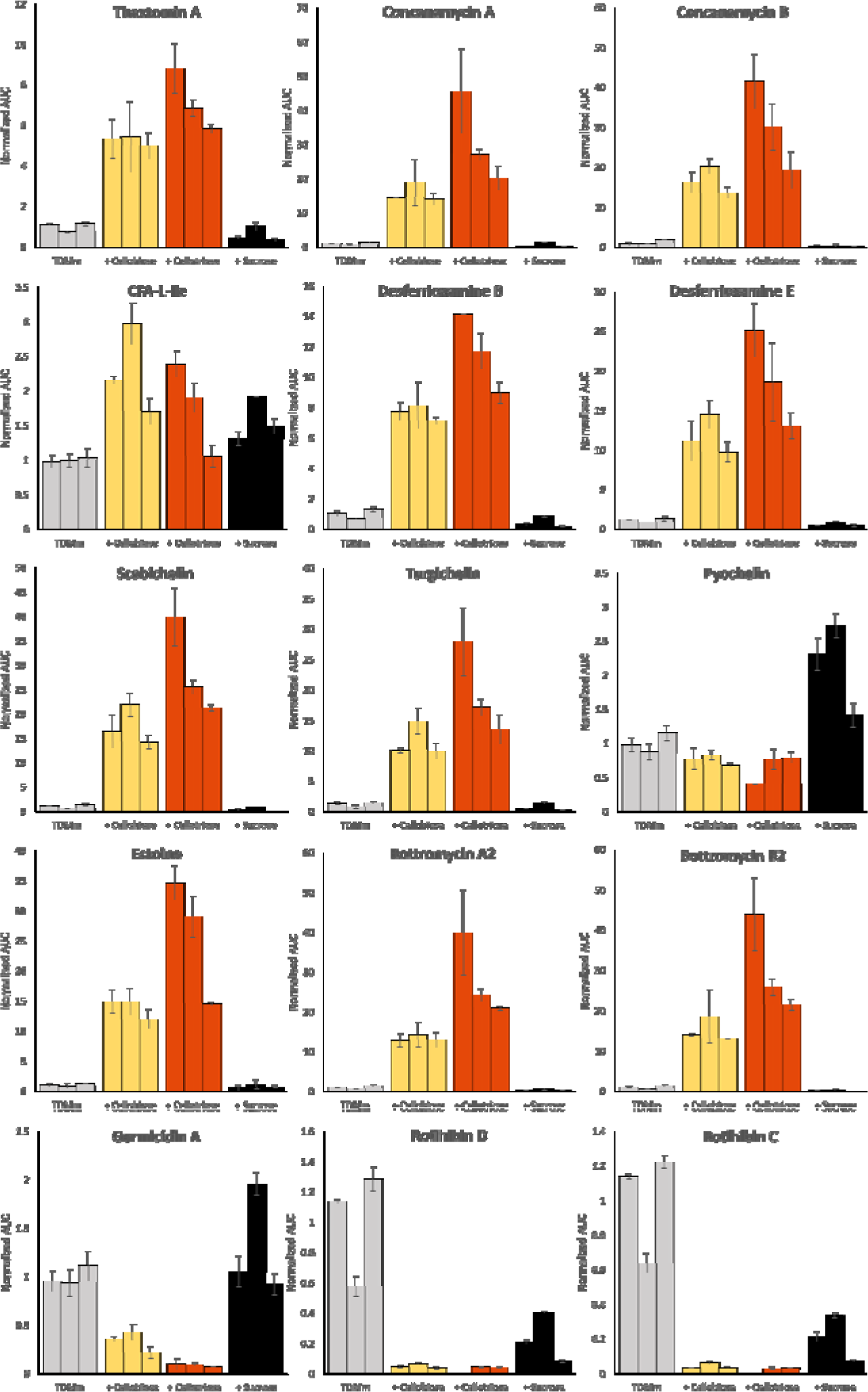
Relative production of the specialized metabolites of *S*. *scabiei* 87-22 upon addition of cello-oligosaccharides. Production levels were assessed in four culture conditions: TDM + maltose 0.5% (TDMm, grey) supplemented with 2.5 mM of cellobiose (+Cellobiose, yellow), cellotriose (+Cellotriose, red) or sucrose (+Sucrose, black). Bar plots display the Area Under the Curve (AUC) of ion peaks normalized to the first replicate of the TDMm condition for each metabolite. Three biological replicates were performed for each culture condition and error bars display the standard deviation observed between three technical replicates.

Regarding the production patterns of the siderophores produced by *S. scabiei* 87-22, both desferrioxamines (B and E) showed enhanced production when *S. scabiei* 87-22 was grown in the presence of cellobiose or cellotriose, with a pattern similar to that observed for thaxtomin A and concanamycins. Desferrioxamine E was the most overproduced of the two, especially in TDMm + cellotriose with an average of 19-fold increase in its abundance levels (Figure 1). Scabichelin and turgichelin – both synthesized by BGC#33a – followed the same trend as desferrioxamines: production increases of about 15- and 25-fold were observed upon addition of cellobiose and cellotriose, respectively. By contrast, pyochelin production was not influenced by either of the cello-oligosaccharides, while sucrose had a limited positive effect of about 2-fold on the siderophore abundance (Figure 1).

The production of bottromycins A2 and B2, as well as its other detected forms (D and E, data not shown), positively responded to the addition of cellobiose and cellotriose (Figure 1). While there was on average a 13-fold overproduction of these antimicrobial metabolites following the addition of cellobiose, cellotriose triggered about twice as much (28-fold increase) production of bottromycins. The production of the osmoprotectant ectoine also positively responded to the presence of cellobiose and cellotriose, with 14- and 26-fold overproduction, respectively (Figure 1).

Out of all the analysed metabolites, only two types of compounds had their relative abundance decreased upon cello-oligosaccharide supply i.e., germicidins and rotihibins. Germicidin A, the inhibitor of *Streptomyces* spore germination, showed a significant decrease in its production levels in both conditions containing cello-oligosaccharides – about 3-fold in TDMm + cellobiose and 10-fold in TDMm + cellotriose (Figure 1). The production of germicidin B displayed the same pattern (data not shown). Similarly, both rotihibins (C and D), recently identified as a novel category of herbicides produced by *S. scabiei* (17), were strongly underproduced with decreases of about 30-fold following cellobiose and cellotriose supply. The addition of sucrose also reduced the amount of rotihibins detected, but to a lower extent (about 5-fold reduction) (Figure 1).

### Tanscriptional response of BGCs to cellobiose and cellotriose

#### BGC of known metabolites under expression control of cello-oligosaccharides

Next to the metabolomic study described above, we also assessed which BGCs responded to the environmental triggers cellobiose and cellotriose at the transcriptional level by RNA-seq. For this, RNA samples were collected 1 and 2 hours post addition of cellobiose and cellotriose (see Methods for details). From all expression data, we focused on the genes belonging to the 18 BGCs involved in the production of known specialized metabolites of *S. scabiei* 87-22 (Table 2, Figure 2). We only considered the expression data of the core biosynthetic gene(s) of each BGC (Table S2) in order to have the best possible correlation between the transcriptomic data and the metabolomic study described earlier. The transcriptional response of the core biosynthetic genes of these “known” BGCs upon supply of cellobiose and cellotriose is displayed in Figures 2A and 2B, respectively, and fold-change values for the core biosynthetic genes of each BGC are displayed in Table 2.

**Figure 2.**
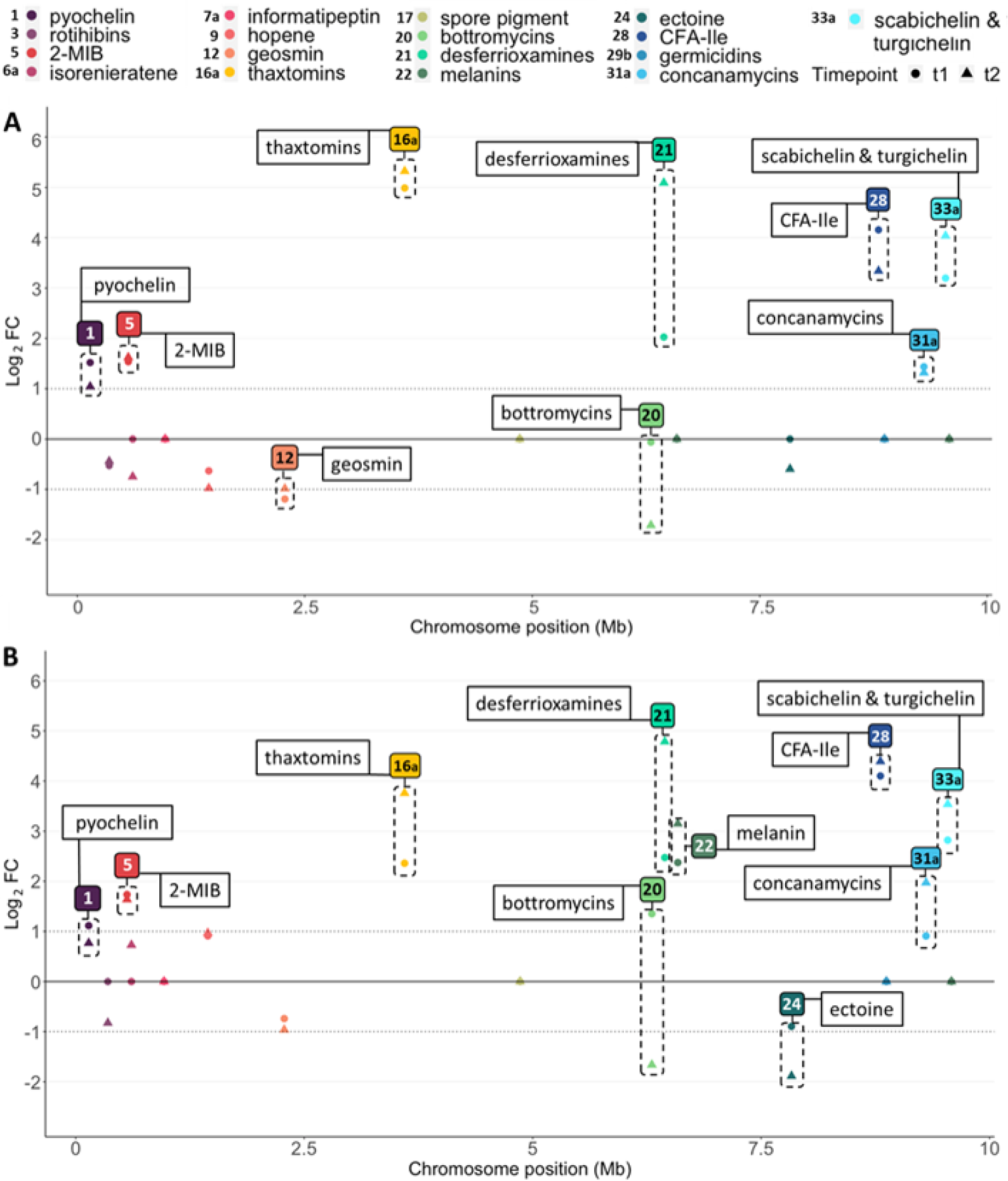
Expression response of core genes of the known BGCs in presence of cellobiose (A) and cellotriose (B). The RNA-seq transcriptomic analysis of *S. scabiei* was conducted in TDM medium (maltose 0.5%) in the presence of cellobiose (A) and cellotriose (B) at time point t1: 1 hour (indicated by circle) and time point t2: 2 hours (indicated by triangle) after induction. The x-axis presents the position of BGCs on the chromosome and the y-axis presents the Log_2_ of the expression fold-change (FC) compared to time point 0 (just before cello- oligosaccharide addition). Only data with significant fold-changes (p-value < 0.05) are displayed (BGCs not meeting this criterium have been set to 0). BGCs with a fold-change above or below the threshold -1 > Log_2_FC > 1 (at least one time point) are highlighted by a dotted frame.

**Table 2.**
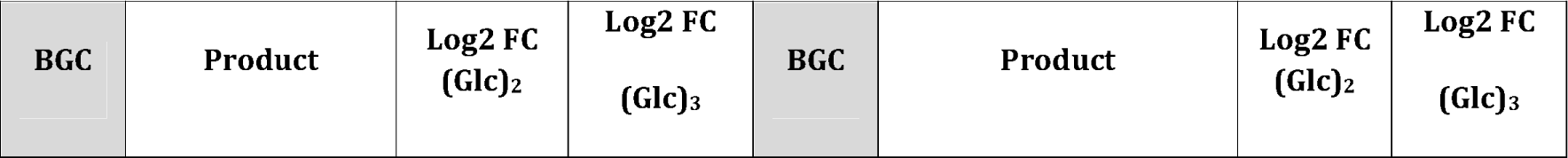
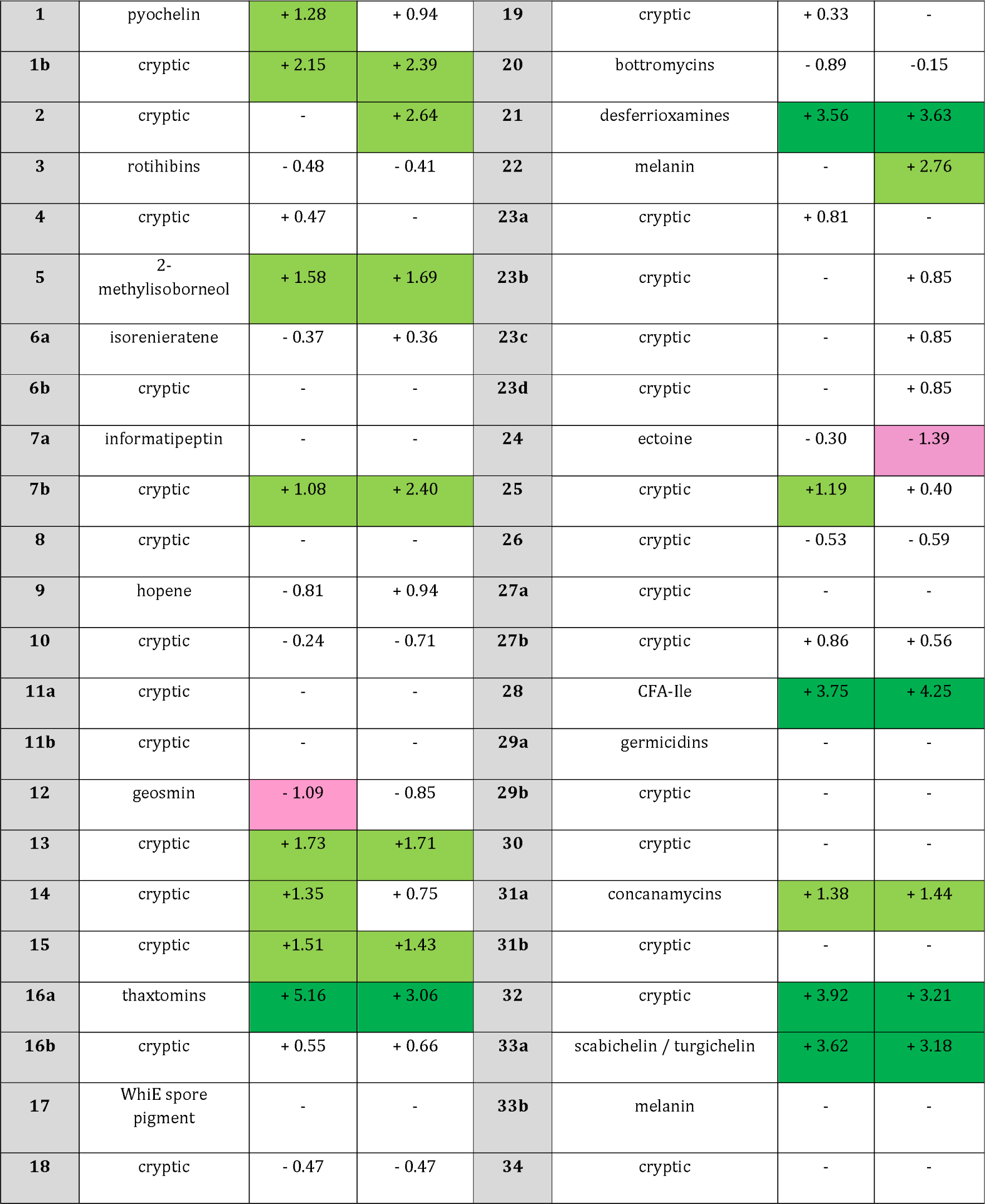
Transcriptional response of BGCs (core genes) upon cellobiose and cellotriose supply

The thaxtomin biosynthesis cluster, designated here as BGC#16a (Table 1), contains two genes whose expression is known to be triggered by both cellobiose and cellotriose (25). Therefore, the response of these core biosynthetic genes, *txtA* (*scab_31791*) and *txtB* (*scab_31781*), can be regarded as “positive controls” for cellobiose/cellotriose upregulated genes. As anticipated, both carbohydrates could drastically increase the transcription of thaxtomin biosynthetic genes. The best transcriptional activation response for *txtA* and *txtB* was observed in the cellobiose condition, i.e., 28- and 56-fold upregulation 2 hours post induction for *txtA* and *txtB*, respectively (Table 2, Figure 2A). Cellotriose was similarly able to activate the expression of both genes, with the biggest fold change also observed at 2 hours post induction, i.e., 10- and 17-fold upregulation for *txtA* and *txtB*, respectively. Analysis of the expression patterns of the other *txt* genes revealed that the whole BGC positively responds to both elicitors, *txtC* displaying the best transcriptional response in the cellobiose condition, with a 143-fold increased expression at 2 hours post induction (Supplementary Figure S1). Overall, the results obtained for the *txt* cluster demonstrate that our experimental set up is appropriate to assess the transcriptional response when *S. scabiei* 87-22 triggers the expression of its main virulence determinant.

Regarding the other BGCs involved in the production of plant-associated molecules, results from the metabolomic and transcriptomic approaches corroborate. As mentioned above, the production of concanamycins was highly responsive to both cello-oligosaccharides. The transcriptomic study confirms this, as the expression of the core biosynthetic genes of BGC#31a, namely *conABCDEF*, showed an average of 2.6- and 3-fold upregulation in cellobiose and in cellotriose, respectively. Genes of BGC#28 responsible for CFA-Ile production also showed a very strong positive expression response to both cello- oligosaccharides (an average of 14- and 19-fold upregulation for cellobiose and cellotriose, respectively), which is in line with the two-fold overproduction of CFA-Ile induced by both cello-oligosaccharides (Table 2, Figure 2). Finally, the analysis of the biosynthetic genes of rotihibins (BGC#3) revealed that the expression of core genes of this cluster does not specifically responds to either cellobiose or cellotriose (Table 2, Figure 2), while the results of our metabolomic analysis suggested a possible down-regulation (Figure 1).

Regarding the biosynthetic genes of desferrioxamines (BGC#21), scabichelin and turgichelin (BGC#33a) siderophores, their expression was also greatly influenced by both saccharides. Transcription of genes responsible for desferrioxamines biosynthesis displayed their greatest response 2 hours post induction for both elicitors, i.e., 40- and 27-fold upregulation for cellobiose and cellotriose, respectively. Biosynthetic genes of BGC#33a involved in scabichelin and turgichelin production were similarly positively affected by cellobiose and cellotriose with an average of 13- and 9-fold upregulation, respectively (Table 2, Figure 2). However, the expression of pyochelin biosynthetic genes (BGC#1a) remained quite steady upon addition of either cello-oligosaccharides, which tends to confirm the metabolomic data (Table 2, Figure 2).

Regarding the BGCs responsible for the production of the antimicrobial compounds bottromycins, and informatipeptin, none of them showed transcriptional activation upon addition of either cellobiose or cellotriose (Table 2). Indeed, the transcription of BGC#20 (for bottromycins) displayed contradicting transcriptional responses (up-regulated or no change at 1 h post induction and down-regulated at 2 h post induction, Figure 2), that overall are not in line with the remarkable overproduction observed via the comparative metabolomic approach (Figure 1, see Discussion). BGC#7a, which shows about 60% synteny to the antimicrobial lanthipeptide informatipeptin, has an expression pattern that was neither influenced by cellobiose nor cellotriose (Table 2, Figure 2). Finally, BGC#29b associated with germicidin production is one of the rare BGCs for which transcriptomic and metabolomic analyses do not correlate, as the absence of transcriptional response is not in line with the marked decrease in production presented in Figure 1 (see Discussion).

Regarding the nine BGCs which belong to the so-called core metabolome of *Streptomyces* species, the addition of cello-oligosaccharides only significantly influenced the expression of BGC#5 (2-methylisoborneol), BGC#12 (geosmin), and BGC#22 (melanin) (Figure 2, Table 2). The effect of both inducers on the expression of the desferrioxamine BGC – which is part of the core metabolome – has already been discussed in the section associated with siderophore BGCs. Cellobiose and cellotriose both activated expression of BGC#5 at almost the same level – around 3-fold, whereas only cellotriose had an impact on the expression of the biosynthetic genes of BGC#22 with an average of 7-fold upregulation. Regarding the osmoprotectant ectoine (BGC#24 in Table 1), we did not observe any significant expression change which does not correlate with the overproduction measured via the metabolomic approach.

#### Cryptic BGCs transcriptionally activated by cello-oligosaccharides

Aside from the 18 BGCs involved in the production of known metabolites, we also assessed the effect of each cello-oligosaccharide on the expression level of the 28 cryptic or orphan BGCs deduced from genome mining (Tables 1 and 2). As shown in Figure 3, the expression of nine cryptic BGCs was influenced by cellobiose or cellotriose. Both carbohydrates significantly increased the transcription of five BGCs, namely BGC#1b (NRPS), BGC#7b (bacteriocin), BGC#13 (bacteriocin), BGC#15 (siderophore), and BGC#32 (siderophore) (Table 2). The highest transcriptional response was observed for the genes of BGC#32 involved in the synthesis of a siderophore metabolite, which were induced up to 28-fold by cellobiose and 15-fold by cellotriose (Table 2, Figure 3). The transcription of another BGC coding for an unknown siderophore (BGC#15) was also increased to about 3-fold by both carbon sources (Table 2, Figure 3). Interestingly, both bacteriocin types of metabolites, i.e., BGC#7b and BGC#13, were upregulated by both cello-oligosaccharides. BGC#13 was equally upregulated by both sugars (around 3-fold), whereas the transcriptional response of BGC#7b was induced more by cellotriose (5.5-fold) compared to cellobiose (2.7-fold). The presence of the cellobiose and cellotriose also positively influenced the transcription of two unknown NRPS BGCs (BGC#1b and BGC#25) with an average 5-fold change for BGC#1b and around 2-fold change for BGC#25 (Table 2, Figure 3).

**Figure 3.**
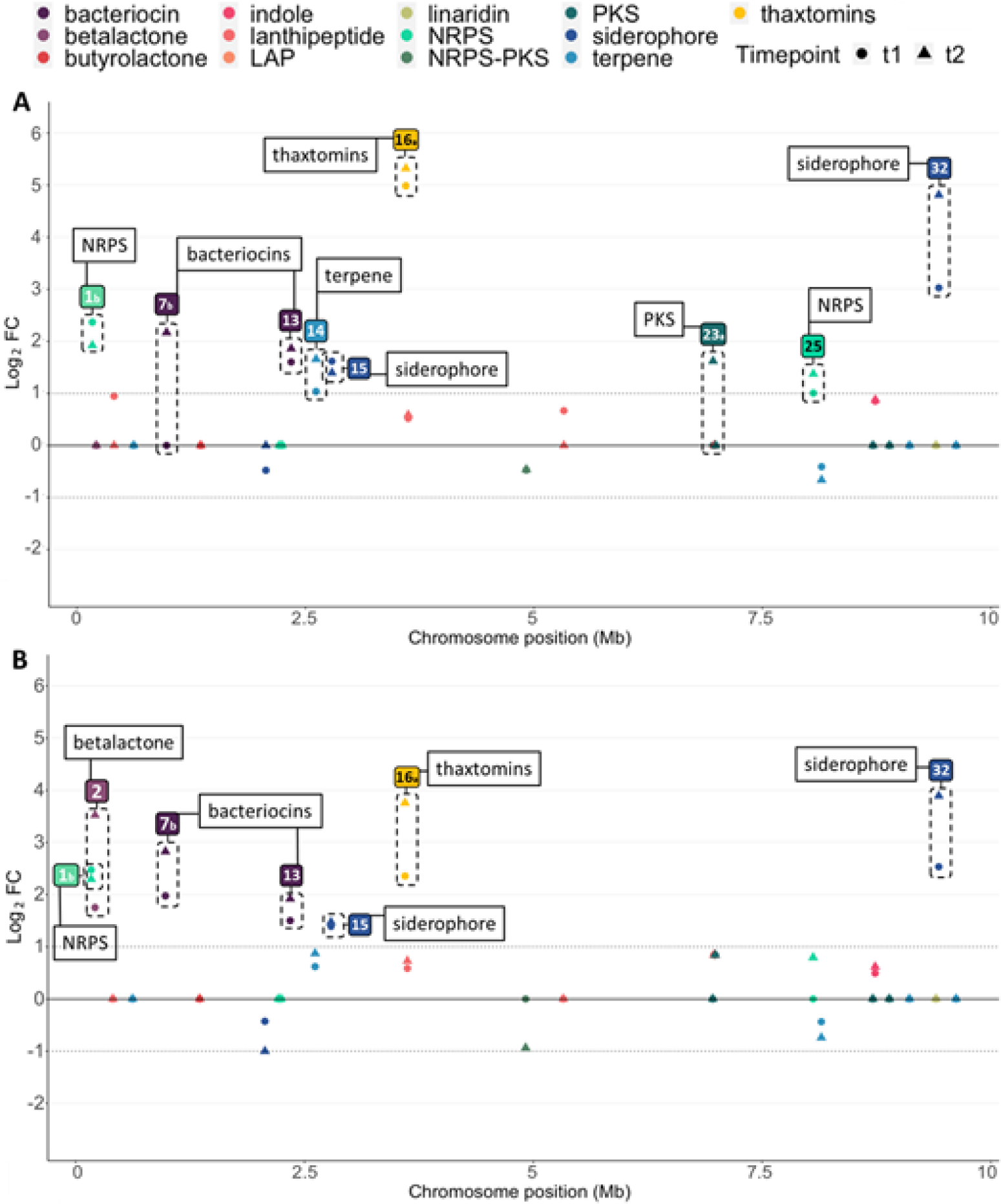
Profile of expression level of core genes of the cryptic BGCs in the presence of cellobiose and cellotriose at two time points. The RNA-seq transcriptomic analysis of *S. scabiei* was conducted in TDM medium (maltose 0.5%) in presence of a) cellobiose and b) cellotriose at time point t1: 1 hour (indicated by circle) and time point t2: 2 hours (indicated by triangle) after induction. The x-axis presents the position of BGCs on the chromosome, whereas the y-axis presents the Log_2_ of the expression fold-change (FC) compared to time point 0 (just before cello- oligosaccharide addition). Only data with significant fold-changes (p-value < 0.05) are displayed (BGCs not meeting this criterium have been set to 0). BGCs with a fold-change above or below the threshold -1 > Log_2_FC > 1 (at least one time point) are highlighted by a dotted frame.

Four other BGCs had their expression upregulated by only one of the two tested cello-oligosaccharides. Cellobiose could in addition induce the expression of BGC#14 (terpene), BGC#23a (PKS), and BGC#25 (NRPS) (Figure 3A), whereas cellotriose positively impacted the expression of BGC#2 (betalactone) (Figure 3B). Importantly, the expression of the core genes of 19 of the 28 cryptic BGCs remained silent or was not significantly influenced by cellobiose or cellotriose.

#### Effect of *cebR* deletion on the expression of cello-oligosaccharide-dependent BGCs

The *txt* cluster responsible for thaxtomin production was previously reported to be under direct control of the cellulose utilization repressor CebR (22). Two CebR-binding sites have been discovered within the *txt* cluster which allows the CebR repressor to switch off the expression of the thaxtomin pathway-specific activator TxtR, in turn resulting in the transcriptional repression of the whole thaxtomin BGC. Binding of cellobiose and/or cellotriose to CebR unlocks the system which allows thaxtomin production. According to our transcriptomic analysis, a total of 16 BGCs have their expression increased by the addition of either cellobiose and/or cellotriose (Table 2) suggesting a possible role of CebR as direct transcriptional repressor of other gene clusters of *S*. *scabiei*. A transcriptome analysis was thus performed in order to assess which BGCs, beyond the txt cluster, would also have their expression under control of CebR. For this, *S*. *scabiei* 87-22 (wild-type) and its Δ*cebR* null mutant (Δ*cebR*) were cultured in ISP2 liquid medium, and RNA samples were collected 3 hours after culture inoculation with fresh mycelium. The volcano plot in Figure 4 shows the relative expression of genes that were determined to be “core biosynthetic genes” of the different BGCs (Table S2). As can be seen in this plot (Figure 4), only one “known” BGC shows an increased expression in the Δ*cebR* mutant, which unsurprisingly corresponds to the thaxtomin biosynthetic cluster (BGC#16a). However, three additional cryptic clusters see their core genes’ expression increased, namely, BGC#14, #16b, and #32 coding for terpene, lanthipeptide, and siderophore specialized metabolites, respectively. In the case of BGC#16b, we can also observe that only one out of its three core genes fall into the upregulated category (“UP” at Figure 4). Interestingly, BGC#32 also responded positively to cellobiose and cellotriose (Table 2) and BGC#14 also showed upregulation upon cellobiose supply (Table 2). However, none of the genes from these cryptic BGCs have been predicted to contain a CebR-binding site (*cbs*) in their upstream region, meaning it is unlikely that their expression would be directly regulated by CebR (see Discussion).

**Figure 4.**
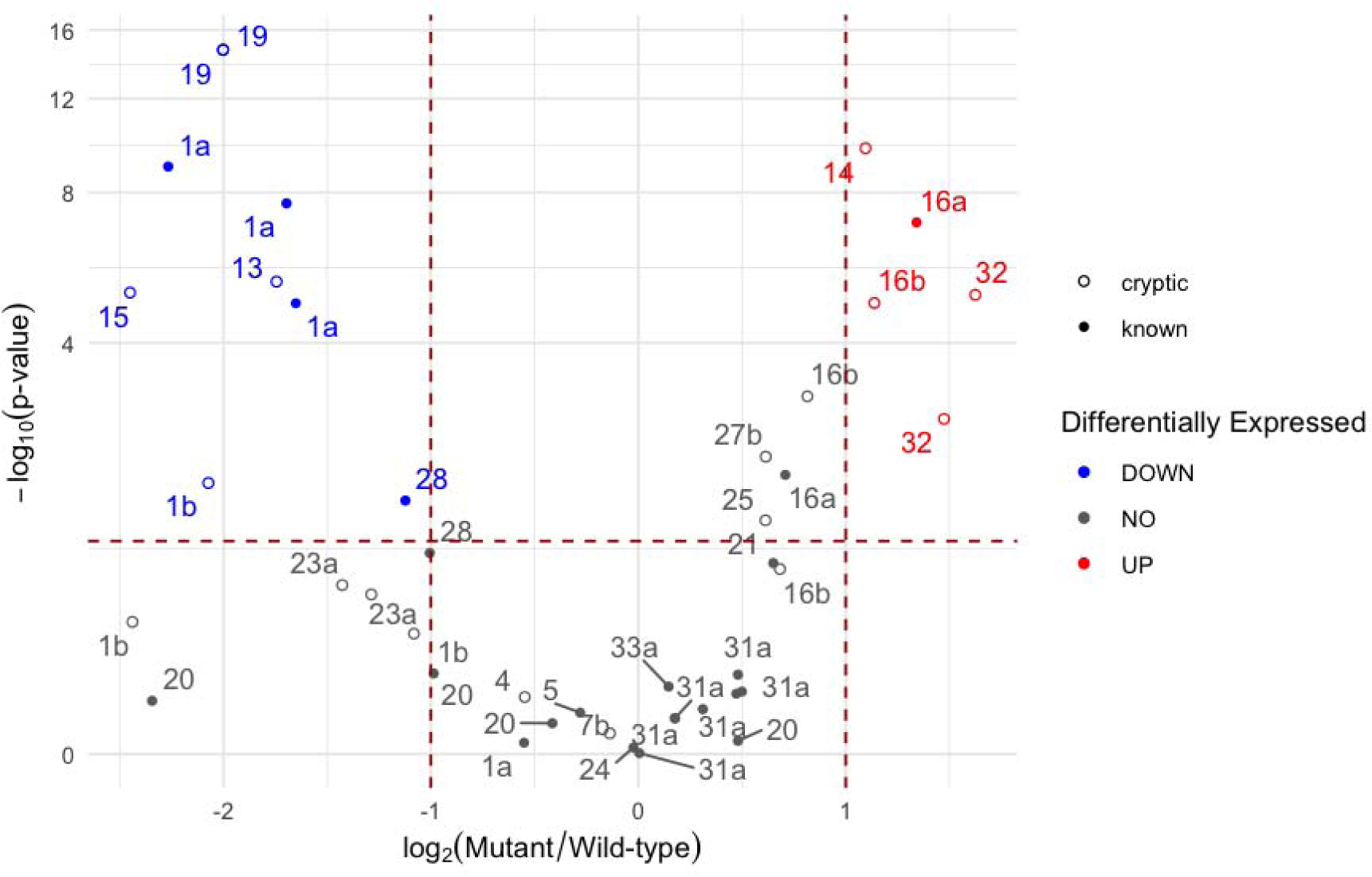
Volcano plot displaying differentially expressed core BGC genes between the *S. scabiei* wild-type strain and the Δ*cebR* mutant. Genes belonging to cryptic BGCs are represented by an empty circle, and those from known BGCs by a full one. Colours indicate the differential expression of each core gene in the Δ*cebR* mutant strain relative to the WT: upregulated (red), downregulated (blue), no significant change (grey). The x-axis displays the Log_2_ fold-change (FC) between the mutant and the WT, while the y-axis corresponds to the -Log_10_ (p-value). Significant expression changes were defined as having a p-value < 0.05 and a Log_2_FC above or below the given threshold, 1 and -1 respectively (-1 > Log_2_FC > 1), these limits are represented by dotted red lines on the plot.

Based again on core gene differential expression, there is a total of 6 downregulated BGCs, among which 2 known clusters (BGC#1a and #28), corresponding to the pyochelin and CFA- Ile biosynthetic clusters. However, these 2 known BGCs showed upregulation upon cellobiose and/or cellotriose supply (Table 2) suggesting there is an absence of correlation between the response to cello-oligosaccharides and the inactivation of the DNA-binding ability of CebR. The four cryptic BGCs that are downregulated correspond to types NRPS (BGC#1b), bacteriocin (BGC#13), siderophore (BGC#15), and lanthipeptide (BGC#19).

## 10. Discussion

### Conclusions on metabolites that respond to cello-oligosaccharides

Of all the different ways to determine what makes an organism excel in a lifestyle or in an environmental niche, generating mutants and assessing their phenotypic repercussions is the most straightforward approach. However, this approach can sometimes lead to erroneous or questionable conclusions for various reasons, such as gene-function redundancy or genetic compensation mechanisms that could lead to phenotypes that understate the importance of a gene. In studies on the CS disease, finding what is really essential for the virulence of *S*. *scabiei* and related species is thus obviously subjected to these constraints linked to the use of reverse genetics. For instance, the inactivation of *scab_1471* in *S. scabiei* resulted in a mutant strain unable to produce pyochelin but showing no sign of reduced virulence, indicating that this siderophore is either not essential for pathogenicity, or that its absence is rescued by other siderophore(s) produced by *S. scabiei* (20). Also, the interference with thaxtomin production caused by the inactivation of the cellobiose and cellotriose beta-glucosidase BglC (41) revealed a genetic compensation phenomenon that awakened the expression of alternative beta-glucosidases allowing *S*. *scabiei* to maintain the capacity to use cello-oligosaccharides (42). The experimental set-ups can also sometimes be suboptimal – such as inappropriate host and/or culture conditions – to observe the real impact of a mutation and therefore to conclude on the role of a gene product in a biological process. This is for example the case of CFA-Ile as gene inactivation in *S*. *scabiei* showed reduced tissue hypertrophy on potato tuber slices, but the impact of the mutant has only been assessed on tobacco *in vivo* and not on its natural hosts which questions the relevance of this molecule in the colonization process (43). Apart from the results on the mutants involved in the biosynthesis of thaxtomins (5,22,23,41), it is thus sometimes difficult to draw conclusions on the importance of a BGC in contributing to the capacity of *S*. *scabiei* to colonize and infect root and tuber plants.

For the above-mentioned reasons, we chose approaches alternative to reverse genetics, i.e., comparative transcriptomic and metabolomic analyses, to determine which part of the specialized metabolism of *S*. *scabiei* would be dedicated to host infection. Both approaches assume that a large proportion of the molecules/genes required for host colonization will respond to the same elicitors and therefore will display production or expression patterns that are synchronized with the main virulence determinants that, in this case, are the thaxtomin phytotoxins. The results of the metabolomic approach provided a clear picture of the specialized metabolites of *S*. *scabiei* that have their production specifically modulated by cello-oligosaccharides. The cello-oligosaccharide-dependent known metabolites of the virulome of *S*. *scabiei* include: i) plant-associated metabolites, namely thaxtomins and concanamycins phytotoxins (and to a lesser extent CFA-Ile), ii) desferrioxamines, scabichelin and turgichelin siderophores, iii) the bottromycin antimicrobials, and iv) the osmoprotectant ectoine. Importantly, of all the known metabolites of *S*. *scabiei*, germicidins, that are autoregulatory inhibitors of *Streptomyces* spore germination, are metabolites that had their production sensibly reduced. Inhibition of germicidin production following the perception of cellotriose that would emanate from the plant cell wall could be regarded as the first “green light” to allow the onset of the pathogenic lifestyle of *S*. *scabiei*. Moreover, the production of the plant growth regulators rotihibins was drastically reduced after addition of cellobiose and cellotriose. This response, opposite to the dynamics of thaxtomins, suggests that rotihibins are not part of the virulome of S. scabiei. The absence of virulence bioassays on natural host plants, the presence of clusters homologous to BGC#3 in plant-helping streptomycetes, and the plant growth promoting effect observed at low doses suggest that rotihibins might be involved in another aspect of the plant-associated lifestyle, despite exhibiting phytotoxicity at higher doses (17).

Our work showed that there is also a strong positive biosynthetic response of desferrioxamines, scabichelin and turgichelin siderophores which are molecules usually produced upon sensing low iron concentrations. Siderophores and iron supply are essential for the onset of both metabolite production and sporulation in streptomycetes (44–46). We have also previously shown that when siderophore biosynthesis responds to signals other than the environmental iron concentration, it can have a strong impact on their developmental program (47). Such a synchronized production of phytotoxins and siderophores, though still being a real enigma regarding the molecular mechanism in place, makes perfectly good sense in terms of host colonization. Iron is mandatory for most housekeeping functions while free iron is not available within the hosts. The upregulation of two additional cryptic BGCs predicted to be involved in the biosynthesis of siderophores (see below) further emphasizes the essential role of iron acquisition during host colonization. Interestingly, Liu et al 2021 reported the absence of production of scabichelin and turgichelin in OBA (19, 28), suggesting that other compounds present in this more complex medium interfere with the elicitor role of cello-oligosaccharides. It has to be noted that production of turgichelin - together with scabichelin - by *S*. *scabiei* (BGC#33a) is reported for the first time in this work. Among the other metabolites whose production differed with the metabolomic analysis performed in OBA (28), we could mention bottromycins which were detected in OBA, but with very little production, while we showed that cello-oligosaccharides strongly induce their production. On the other hand, compounds that were reportedly produced to high levels in OBA – such as concanamycins, thaxtomins and desferrioxamines – were also detected and overproduced upon cello-oligosaccharides addition. Finally, in their metabolomic analysis, Liu and colleagues (28) also highlighted the abundance of CFA-Ile in OBA culture extracts (also shown in (43)). The presence of CFA-Ile in our culture extracts indicates an important role for this molecule in the virulome, even though the addition of cello-oligosaccharides only has a limited positive impact on CFA-Ile production (Figure 1), while we observed massive overexpression via our transcriptomic analysis (Figure 2 and Table 2). Surprisingly, we previously reported in a proteomic study that the abundance of two proteins of the CFA-Ile biosynthetic pathway – SCAB79611 (Cfa2) and SCAB79671 (CFL) – significantly decreased upon cellobiose addition (48). Altogether, these results suggest that the production levels and the expression response of CFA-Ile biosynthetic proteins/genes are highly sensitive to the chosen culture conditions which could explain the important differences between our experimental setup conducted here in minimal media and earlier studies.

Additional metabolites – identified in a previous study (28) – were found in all the tested culture conditions. Production of aerugine, andrachcinidine, Cyclo(L-Val-L-Pro), and a form of the plant hormone auxin (Indole-3-acetic acid - IAA) was more or less influenced by the addition of cello-oligosaccharides, but not dramatically (Figure S2). Only andrachcinidine, an alkaloid metabolite putatively involved in plant defence, was induced by cellobiose and the production of IAA was reduced by a factor two in TDMm + cellobiose and cellotriose. Further research is required to link these metabolites with their currently cryptic BGCs, except for IAA whose biosynthetic genes have previously been identified (49). On the other hand, the presence of additional metabolites described by Liu and colleagues (28) was investigated, but none of them was found in our extracts. These metabolites were: mairine B, bisucaberin, dehydroxynocardamines, and 211 A decahydroquinoline. Informatipeptin – a bioactive compound associated with antimicrobial activity – most-likely produced by BGC#7a could not be detected in any of our extracts.

### Cellobiose versus cellotriose as elicitors of virulence

Another question we wanted to address through this work is whether cellotriose could, equally to cellobiose, trigger the “virulome” of *S*. *scabiei*. Indeed, if most studies have been performed with cellobiose as elicitor - the product being much less expensive and available in larger quantities compared to cellotriose -, earlier works suggest that instead, cellotriose is more likely to emanate from living plants considering the cell wall-related action of thaxtomins. Indeed, cellotriose was shown to be naturally released from actively growing plant tissue, and a treatment with pure thaxtomin A increased the amount of cellotriose exuded by radish seedlings (25). On the other hand, we proposed that cellobiose would rather result from cellulolytic degradation of dead plant material (27). Our metabolomic and transcriptomic analyses revealed that cellobiose and cellotriose are able to induce a similar response. It is however important to note that for most metabolites of the virulome, cellotriose was a stronger inducer compared to cellobiose (Figure 1). Even for germicidins, one of the two metabolites whose production is reduced and not increased by cello- oligosaccharides, cellotriose had a stronger impact. One possible explanation is that, once internalized, cellotriose is hydrolysed to cellobiose and glucose by the beta-glucosidase BglC, therefore further providing the disaccharide eliciting molecule in the intracellular compartment. However, once inside the cell, the hydrolysis of cellobiose by BglC generates two molecules of glucose that will feed glycolysis (primary metabolism) and no longer act as trigger for the specialized metabolism of *S*. *scabiei*. In contrast, our transcriptomic analysis suggested that cellobiose was in general a better elicitor compared to cellotriose (Figure 2 and 3). This is not surprising as we showed that the import of cellobiose is faster than that of cellotriose in *S*. *scabiei* (41), which explains the observed slower transcriptional response. Also, RNA samples were collected after 1 h and 2 h post addition of either cello-oligosaccharides while the extracts for the metabolomic analysis were collected after 96 h of growth. The short-term transcriptional response thus cannot be quantitatively compared to the long-term metabolite production response.

The most important observation is that, for the large majority of BGCs and known metabolites that have been investigated here, we saw a clear correlation between the data obtained via the transcriptomic and metabolomic approaches. Among the few exceptions we can mention the case of the osmoprotectant ectoine, its production being highly induced by both cello-oligosaccharides (Figure 1) while we could not see significant expression changes (Figure 2 and Table 2). The most plausible explanation lies in the fundamental difference in the culture conditions as samples for RNA-seq analyses were collected from liquid cultures just after the addition of the elicitors while metabolite samples were extracted from 96 hours solid cultures. Osmotic protection is expected to be more important after 96-hours cultures at the agar-air interface compared to a couple of hours in liquid cultures with no osmotic changes. The expression of genes involved in ectoine biosynthesis could also be controlled by development-related signals or regulators that are not yet available at RNA sampling time points. Also, no correlation was observed between the strongly reduced germicidin production (Figure 1) and the negligible transcriptional response of the corresponding BGC#29a (Figure 2 and Table 2). Again, the lack of expression change could be explained by the differences in culture conditions. Finally, the transcriptional response of the BGC#20 for bottromycins was too irregular (up-, down-regulation, or no changes according to the elicitors and time points) to make any correlation with the strong overproduction observed via the metabolomic study.

### CebR-independent response of most cello-oligosaccharide-dependent BGCs

Surprisingly, except for the thaxtomin gene cluster, only 2 of the 15 other BGCs that showed a strong transcriptional response to cellobiose and cellotriose also showed overexpression in the Δ*cebR* mutant (Figure 4). This result, though unexpected, is in line with the absence of CebR-binding sites in the upstream region of pathway-specific transcriptional activators and core biosynthetic genes of these cello-oligosaccharide expression-dependent BGCs. Through our earlier proteomic analysis, we also observed that many proteins whose production was activated in *S*. *scabiei* 87-22 by cellobiose did not show production changes in the *cebR* null mutant (48). This could be explained by the possible ability of CebR to bind to “non-canonical” DNA sequences, as previously observed for other transcription factors that also link nutrient sensing and the specialized metabolism (50). Alternatively, cellobiose and cellotriose could be sensed by another and yet unknown transcription factor.

### Cryptic metabolites and perspectives

Although strain *S*. *scabiei* 87-22 is rather well-studied as a model organism, the plurality of its cryptic and/or silent BGCs highlighted a huge reservoir of yet unknown metabolites. Nine of these cryptic BGCs showed a significant response to either both or one of the two cellooligosaccharides, suggesting that some of these unknown compounds may also be part of the virulome of *S*. *scabiei*. The best transcriptional response was observed for BGC#32 (Table 2) involved in the synthesis of a siderophore type metabolite. Together with the transcriptional awakening of BGC#15, the cello-oligosaccharide-dependent response of siderophore-related BGCs further underlines the importance of iron acquisition during host colonization. We also observed the positive expression changes for two BGCs responsible for the production of two bacteriocin type metabolites (BGC#7b and #13). The strain-specificity of these antibacterial peptides is unknown but their synchronized biosynthesis with other host colonization molecules could be seen as a strategy to prevent competing soil-dwelling bacteria to also access the starch reservoir of tubers.

The structure and the bioactivity of the metabolites whose production is triggered by cellobiose and cellotriose is currently under investigation and should lead to the identification of new key virulence determinants associated with the common scab disease.

### 11. Author statements

### 11.1 Authors and contributors

Benoit Deflandre: Conceptualization, Formal analysis, Investigation, Methodology, Supervision, Visualization, Writing – original draft

Nudzejma Stulanovic: Formal analysis, Investigation, Software, Visualization, Writing – original draft

Sören Planckaert: Formal analysis, Investigation, Methodology, Writing – review & editing

Sinaeda Anderssen: Data curation, Formal analysis, Methodology, Software, Visualization, Writing - review & editing

Beatrice Bonometti: Investigation Latifa Karim: Investigation, Methodology, Resources Wouter Coppieters: Methodology, Project administration, Resources, Supervision Bart Devreese: Funding acquisition, Methodology, Project administration, Supervision, Writing – review & editing Sébastien Rigali: Conceptualization, Funding acquisition, Methodology, Project administration, Supervision, Visualization, Writing – original draft

### 11.2 Conflicts of interest

The authors declare that there are no conflicts of interest

### 11.3 Funding information

The work of S.R. and Be.D was supported by an Aspirant grants from the FNRS (grant 1.A618.18) and FRIA grants from the FNRS for S.R. and S.A. (FRIA 1.E.031.18-20) and S.R. and N.S. (FRIA 1.E.116.21). Ba.D. and S.P. were also supported by a Bijzonder Onderzoeksfonds (BOF, grant 01B08915)-basic equipment from the Ghent University special research funds. S.R. is a Fonds de la Recherche Scientifique (FRS-FNRS) senior research associate.

### 11.4 Ethical approval

Not applicable

### 11.5 Consent for publication

Not applicable

## Acknowledgements

[An Acknowledgements section is not compulsory. However, if materials and results were obtained from outside the authors’ laboratories (e.g. production of antibodies, properties of strains), this must be acknowledged.]

---

**Supplementary Figure S1.**
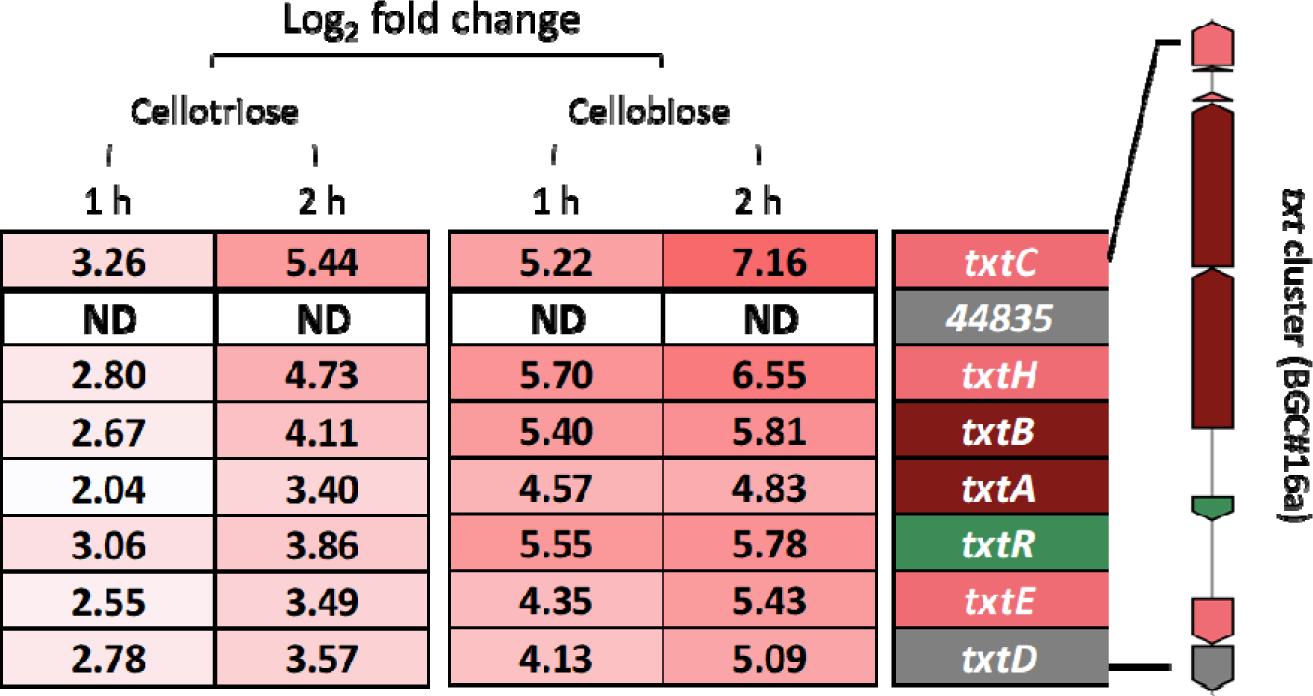
Detailed expression fold changes (expressed as Log_2_) for individual genes of the *txt* cluster (BGC#16a) in cellobiose and cellotriose after 1 or 2 hours of incubation. ND = Not Detected

**Supplementary Figure S1.**
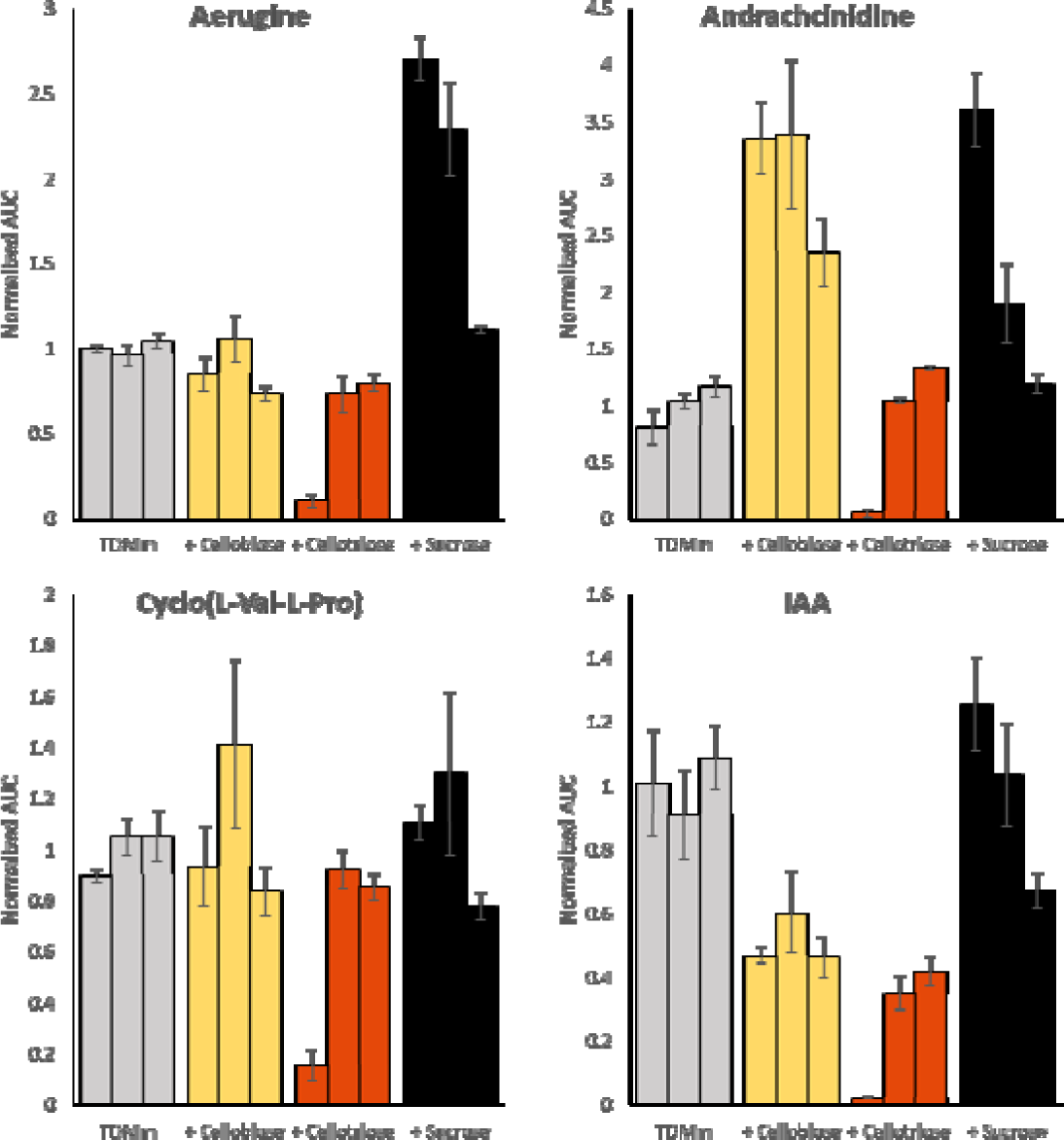
Relative production of the specialized metabolites of *S*. *scabiei* 87-22 upon addition of cello-oligosaccharides. Production levels were assessed in four culture conditions: TDM + maltose 0.5% (TDMm, grey) supplemented with 2.5 mM of cellobiose (+Cellobiose, yellow), cellotriose (+Cellotriose, red) or sucrose (+Sucrose, black). Bar plots display the Area Under the Curve (AUC) of ion peaks normalized to the first replicate of the TDMm condition for each metabolite. Three biological replicates were performed for each culture condition and error bars display the standard deviation observed between three technical replicates.

**Supplementary Table S1.**
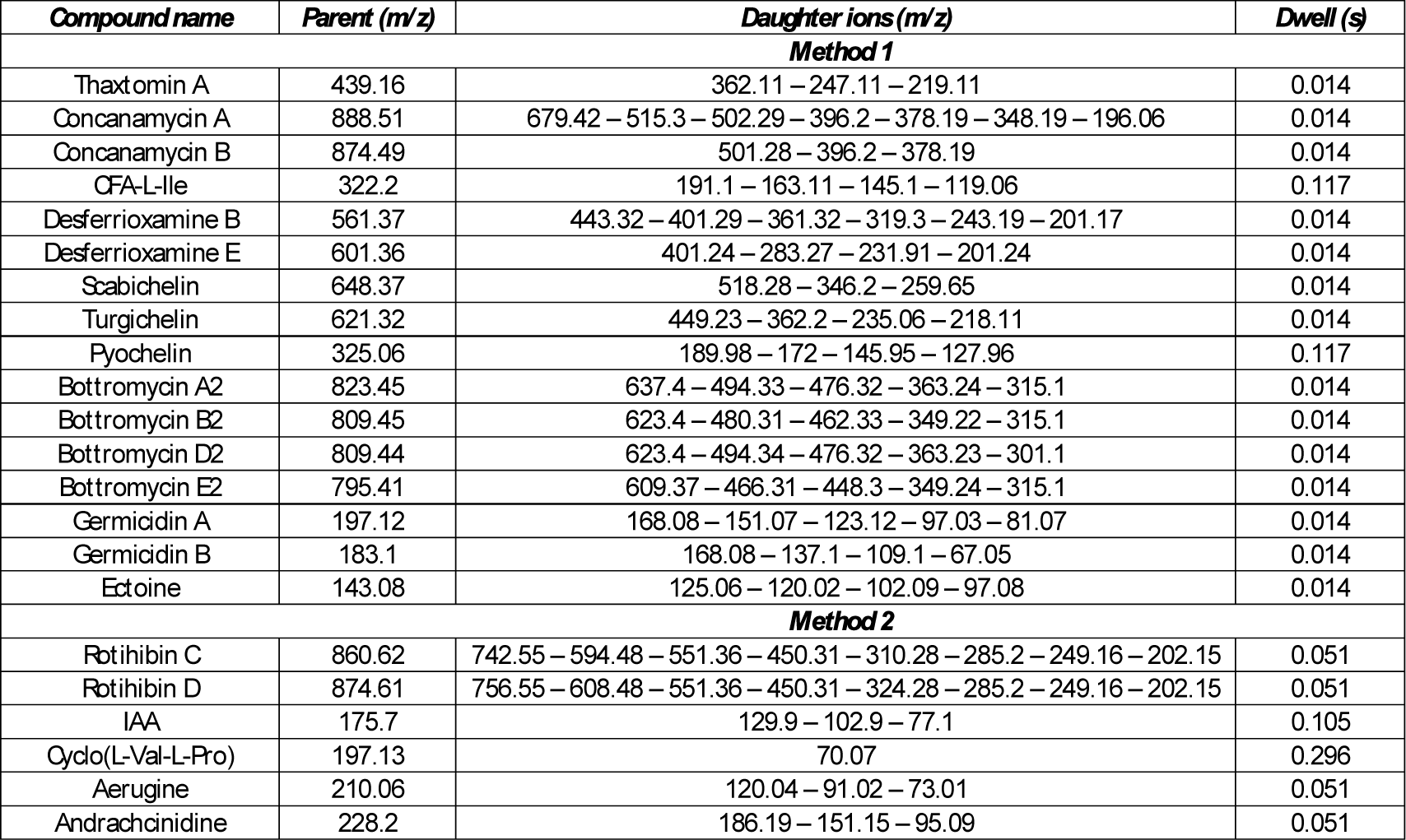
The MS conditions for the metabolites of interest in MRM mode.

**Supplementary Table S2.**
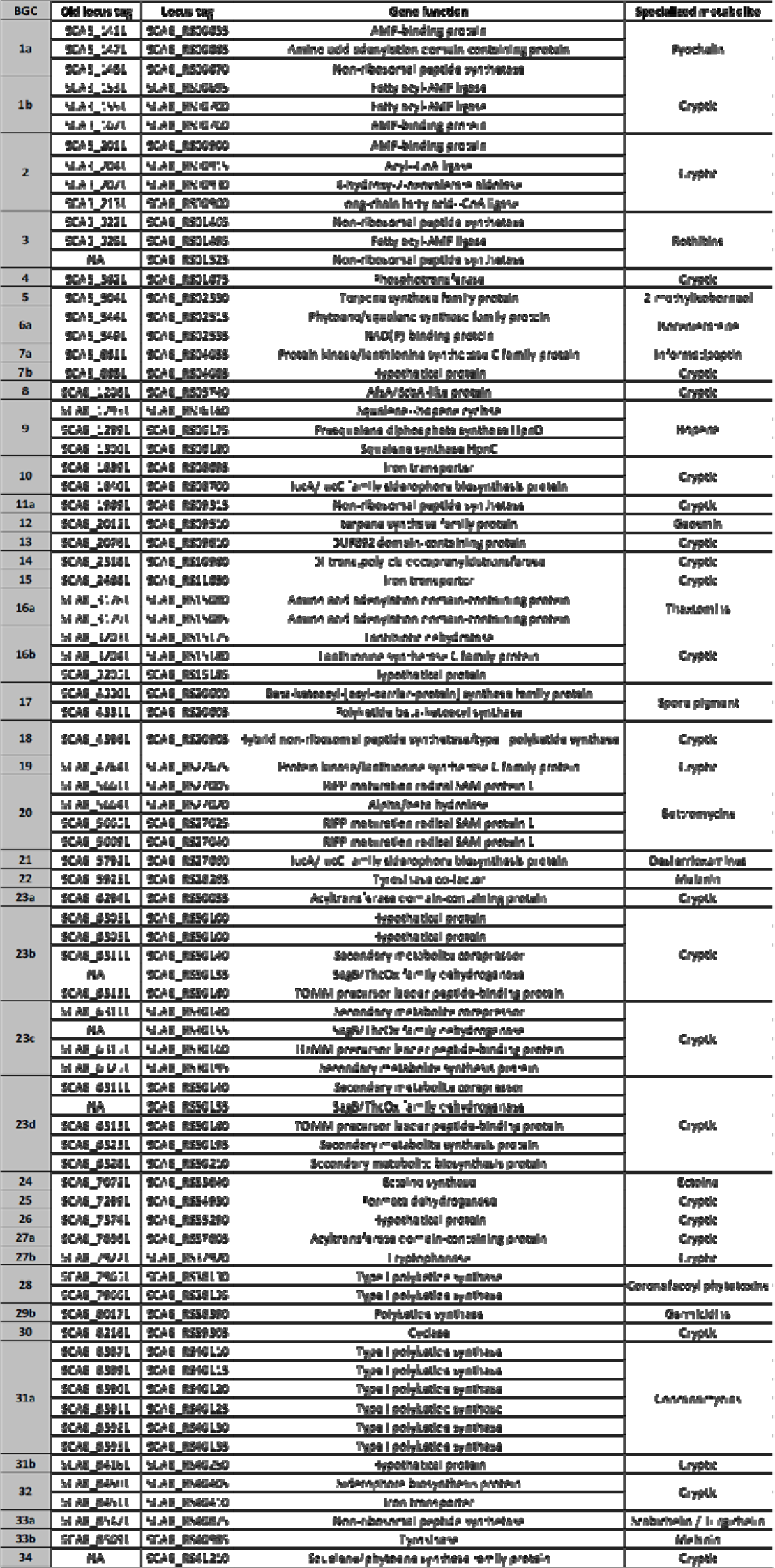
List of the core biosynthetic gene(s) and their predicted function for each BGC

## Notes

### Competing Interest Statement

The authors have declared no competing interest.

